# SRST2: Rapid genomic surveillance for public health and hospital microbiology labs

**DOI:** 10.1101/006627

**Authors:** Michael Inouye, Harriet Dashnow, Lesley Raven, Mark B. Schultz, Bernard J. Pope, Takehiro Tomita, Justin Zobel, Kathryn E. Holt

## Abstract

Rapid molecular typing of bacterial pathogens is critical for public health epidemiology, surveillance and infection control, yet routine use of whole genome sequencing (WGS) for these purposes poses significant challenges. Here we present SRST2, a read mapping-based tool for fast and accurate detection of genes, alleles and multi-locus sequence types (MLST) from WGS data. Using >900 genomes from common pathogens, we show SRST2 is highly accurate and outperforms assembly-based methods in terms of both gene detection and allele assignment. Here we have demonstrated the use of SRST2 for microbial genome surveillance in a variety of public health and hospital settings. In the face of rising threats of antimicrobial resistance and emerging virulence amongst bacterial pathogens, SRST2 represents a powerful tool for rapidly extracting clinically useful information from raw WGS data. Source code is available from http://katholt.github.io/srst2/.

---

Rapid molecular typing of bacterial pathogens is critical for public health epidemiology, surveillance and infection control [1, 2]. Two key goals of such activities are: (i) to detect the presence of genes linked to clinically relevant phenotypes - including virulence genes, antimicrobial resistance genes or serotype determinants; and (ii) to classify isolates into clonal groups, via multi-locus sequence typing (MLST [3]) or detection of clone-specific or other epidemiological markers. Whole genome sequencing (WGS) or “genomic epidemiology” is increasingly being adopted for these tasks and has the potential to replace current techniques which are mainly based on PCR and/or restriction enzyme digestion coupled with sequencing or size separation via electrophoresis [1, 4]. WGS is particularly attractive as (i) it can be applied simultaneously to large numbers of bacterial isolates of any species with no need for organism- or target-specific reagents, and (ii) the resulting data is readily shareable, can be compared easily with past and future data sets, and is informative for both routine surveillance (monitoring genes and clones) and detailed outbreak investigation (genome-wide phylogenies for transmission analysis) [2, 4].

WGS has revolutionised pathogen research, and its potential to revolutionise the practice of public health epidemiology, surveillance and infection control has been recognised for some time [4–10]. Despite the enthusiasm and several demonstration studies [11–16], the routine use of WGS poses significant challenges for public health and diagnostic laboratories, foremost of which is a lack of solutions for the rapid and reproducible extraction of informative, interpretable and shareable data from raw sequence data [1, 17].

Currently available methods rely on assembling short reads into longer contiguous sequences (contigs), which can be interrogated using BLAST or other search algorithms to identify genes or alleles of interest (e.g. ARG-Annot [18]; ResFinder, PlasmidFinder and MLST typer [19–21]; BIGSdb [22, 23]). The reliance on assembly introduces efficiency and sensitivity problems due to the data, time and computational requirements for generating high quality assemblies of bacterial genomes from short reads. There are several assemblers (e.g. *Velvet* [24], *SPAdes* [25]) that can produce a bacterial genome assembly in minutes to hours with a few gigabytes of memory. However the production of high quality assemblies with these tools requires quality filtering and other pre-processing of reads, and optimisation of kmer length and other parameters which in practice requires several alternative assemblies to be generated and compared [26, 27], thus multiplying by an order of magnitude the amount of computational time and memory required to produce each genome prior to typing analysis. Further, the quality of even highly optimised assemblies remains highly variable, even for closely related genomes sequenced together in multiplex. Hence assembly-based analyses of genomes sequenced with short-read technology are very difficult to standardize and quality-control, which is important to ensure robust, reliable and reproducible assays for use in public health and infection control.

Here we describe a new tool for genomic epidemiology, SRST2, which performs fast and accurate detection of genes and alleles direct from WGS short sequencing reads. SRST2 can type reads using any sequence database(s) and can calculate combinatorial sequence types defined in MLST-style databases [3]. We demonstrate its utility for routine molecular typing in public health and hospital laboratories via automated MLST and typing of virulence, antimicrobial resistance and plasmid genes. SRST2 is named after our earlier tool SRST (Short Read Sequencing Typing) which performed MLST on short reads [28], however the SRST2 code is entirely novel and uses different read mapping, scoring and reporting algorithms than SRST, is more stable and robust, and is designed for gene detection and allele typing as well as MLST.

## Approach and Implementation

Given a read set and database of reference allele sequences, SRST2 is designed to perform two key tasks: (i) detect the presence of a gene or locus, and (ii) determine the precise or closest matching allele for that locus, amongst a set of possible reference allele sequences. The approach is illustrated in Figure 1. A database of reference sequences must be provided in fasta format, in which the fasta headers indicate both the locus (so that alleles of the same locus can be compared) and a unique name for each allele. In the case of MLST data an additional database of ST profiles is provided as tab-delimited text, which assigns STs to unique combinations of alleles. Current MLST data (allele sequences and profile definitions), suitable for use with SRST2, can be downloaded from pubmlst.org automatically using the *getmlst.py* script supplied with SRST2. Other sequence databases can be easily formatted for use with SRST2 using the scripts supplied with the program. Any number of sequence databases can be analysed in a single run, allowing for simultaneous typing of MLST, resistance genes and virulence genes.

**Figure 1.**
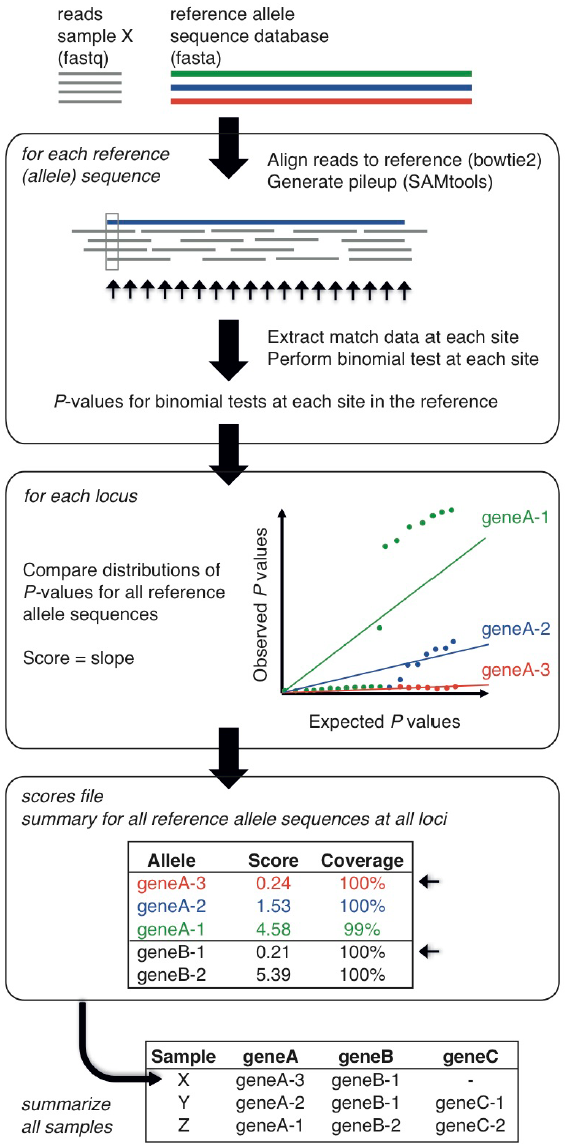
Summary of SRST2 approach. Inputs are reads (fastq format) and one or more databases of reference allele sequences for typing (fasta format). Reads are aligned to all reference sequences (using *bowtie2*) and each alignment processed (using *SAMtools*). At each position in each alignment, the number of matching and mismatching bases is determined and a binomial test is performed to assess the evidence against the reference allele; resulting in a set of P-values for each reference allele sequence. To determine which of all known reference alleles is most likely present at a given locus, the P-value distributions for known alleles are compared as described in the text. Briefly, for each allele the P-values expected if the reads were derived from the reference allele in the presence of a given level of sequencing error (set to 1% of bases by default) are regressed on those actually observed, similar to a Q-Q plot; the slope of the fitted line, which increases with the strength of evidence against the reference allele, is calculated and taken as the score for that allele. The scores file (optional output) contains the scores for each allele at each locus, along with additional information about the alignments for each allele including percent coverage. For each locus, the allele with the lowest score is accepted as the closest matching allele (small arrows) and reported in the output table. In MLST mode, sequence type (ST) definitions are provided as input and used by SRST2 to calculate STs for each read set.

For each input database, reads are aligned using *bowtie2* v2.1.0 or above with the ‘--very-sensitive-local’ and ‘-a’ settings, and all alignments are reported to a file in SAM format. Mapping sensitivity can be fine-tuned by specifying to SRST2 any of the parameters available within the *bowtie2-align* command or a maximum number of mismatches per read (default 10 mismatches allowed). Flags in the resulting SAM file are modified so that each read is included in the pileup for every allele to which it is aligned. Pileups are generated using *SAMtools* v0.1.18 *mpileup* and parsed by SRST2 to determine percent coverage, divergence, and mismatches, and to calculate a score for each possible allele.

### Allele scoring

An overview of the scoring approach is given in Figure 1. We begin with an alignment of reads from sample *s* to a reference sequence *r*. At each position *i* in the reference sequence *r* (*r_i_*), let *s_i_* be the set of reads in sample *s* that align to *r_i_*. Let *a_i_* be the total number of reads in *s_i_*, and let *b_i_* be the number of reads in *s_i_* in which the aligned base does not match the reference base at *r_i_*. If sample *s* contains the precise sequence *r*, then the probability of a mismatched base at any position in an aligned read is equal to the per-base error rate of the sequencing technology *e_i_*, which for Illumina is taken to be 0.01, although this can vary depending on what pre-processing steps are implemented [29, 30].

To quantify the evidence against the presence of the reference sequence *r* in *s*, we perform a Binomial test at each position *r_i_*, to generate a 1-sided P-value *P_i_* to assess the probability of observing *a_i_*-*b_i_* successes in *a_i_* trials, with a probability of success of 1-*e_i_*. Any change at position *r_i_* - including a base substitution, an insertion of any size or a deleted base - is treated as a mismatch, incrementing *b_i_* by 1. For large deletions that result in an absence of any aligned reads (including truncations of the end of the sequence), *a_i_* = 0 and no Binomial test is possible. In this case, the evidence for the deletion is provided by the reads which align adjacent to the deletion but do not align across the deletion. Hence we calculate the average number of reads aligned to the two bases preceding the deletion, *d_i_*, and conduct the Binomial test with *a_i_* = *b_i_* = *d_i_*.

We then utilize a non-parametric approach to score each allele by considering the set of all P-values calculated for reference sequence *r*. First, to minimise artefacts associated with fluctuation in read depths, we (a) set *P_i_*=1 where *b_i_*=0, and weight *P_i_* by the relative read depth (i.e. weight of evidence) at position *r_i_* compared to those of other positions in *r*:

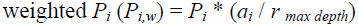

We then compare the sorted −log_10_(*P_i_*) values versus those of the theoretical distribution of −log_10_(*x_j_*/n) where n = length(*r*) and *x_j_ =* 1,2,…n, analogous to inspecting a quantile-quantile (QQ) plot (Figure 1). A linear model is fitted to the two probability distributions and the resulting slope is taken as the score for reference sequence *r*, *score_r_*. Here we leverage a common criticism of linear models to our advantage: the susceptibility to outliers at the tails of the distribution. In this case, outliers are typically SNPs or indels relative to the sequence *r* which, because they result in low P-values in the Binomial test and thus very high values of −log_10_(P), are at the end of the observed distribution (Figure 1). Thus when a linear model is fitted, its slope increases with the number of well-supported SNPs and indels compared to the reference. As a result, among reference alleles of the same locus, the sequence *r* with the lowest *score_r_* (flattest slope in the QQ plot) is the most likely match for sample *s*.

### Reporting outputs

SRST2 output tables report, for each sample *s* and each locus or gene cluster, the lowest scoring allele sequence *r*, the average read depth of *s* across *r* and indicators of any evidence against a precise match with *r* (including mismatches supported by >50% of aligned reads, or read depth falling below a cutoff). Only matches passing the user-set coverage and divergence cut-offs (by default, >90% coverage and <10% divergence) are reported. For MLST data, STs are calculated according to the MLST profiles database provided, based on the closest matching alleles at each locus.

Normally, an exact match between *r* and *s* would be assigned if (a) *r* has the lowest *score_r_* amongst the set of alleles of the same locus or gene cluster, and (b) there are no SNPs or indels between *r* and *s*. If (a) holds but (b) does not, this is indicative of a novel allele and SRST2 will flag the result in output tables. In such cases, we recommend that users who are interested in defining novel alleles should inspect the raw sequence data (which may be assisted by the alignments, pileups and consensus fastq files generated by SRST2).

Optionally, SRST2 can report the full details of scoring *s* against all reference sequences *r*, to enable users to parse and interpret the results to suit specific needs. These include average depth of *s* across *r*, average depth across the first and last two bases of *r*, the number of positions in *r* in which the majority of aligned reads in *s* show a mismatch against *r* (with SNPs, insertion/deletions and truncations reported separately), the depth of bases neighbouring truncations and, for the position with the greatest proportion of mismatching reads, the total aligned reads, total mismatching, proportion mismatching, and Binomial p-value.

### Availability

SRST2 Python code is freely available [31] and utilises *bowtie2* [32] for read mapping and *SAMtools*[33] for alignment processing.

## Validation using real and simulated data

### Validation of allele calling

To assess the accuracy of allele identification with SRST2, we analysed publicly available Illumina data from 543 bacterial genomes of five different species for which independent MLST data was available (Table 1). With seven loci in each MLST scheme, this yielded 3,801 allele calls across 35 loci to assess call rate and false positive rate. The read sets represented a wide range of average read depths, with 90% in the range 12x - 130x and 50% between 20x - 60x (Table 1). For each species, we used SRST2 to download the latest MLST database from pubmlst.org and subsequently ran SRST2 using default parameters. Median run time was 6 minutes per sample (interquartile range, 4-10 minutes) and increased linearly with number of reads (Figure 2). Efficiency can be easily improved or standardised, without data pre-processing, by instructing SRST2 to map the first N reads only.

**Figure 2.**
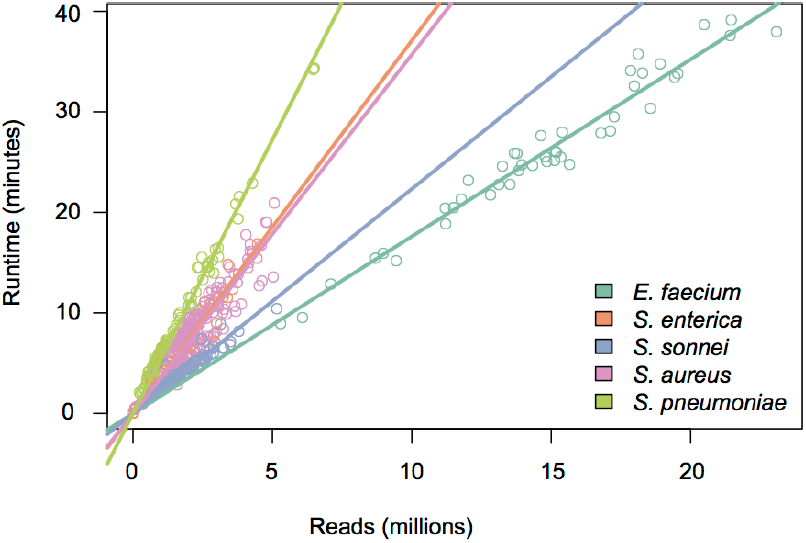
Run times for MLST analysis with SRST2. Lines are linear regression of runtime on reads, calculated separately for each species from public datasets (details in Table 1).

**Table 1.**
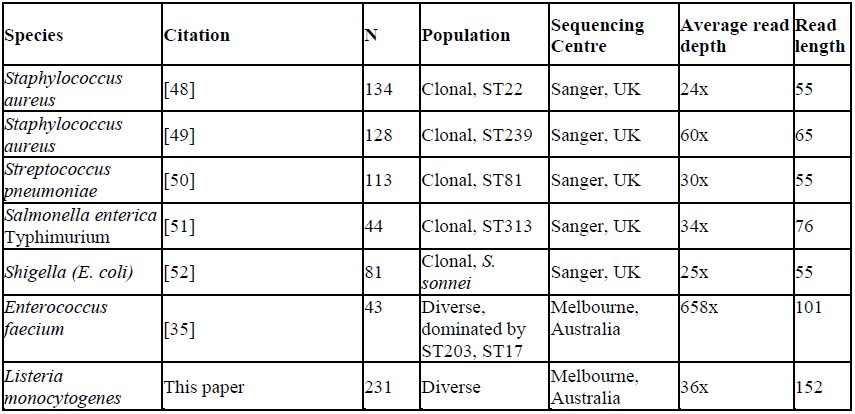
Data sets used to assess accuracy of SRST2

SRST2 call rates and true positive rates increased with average read depth, stabilizing with depths ≥15x (Figure 3a, **Additional file 1**). For comparison, we also assembled each read set using *Velvet* [24] and *VelvetOptimiser* [34] and used nucleotide BLAST to identify MLST alleles (assembly+BLAST method; **Methods**). At read depths ≥15x, SRST2 made significantly more allele calls than assembly+BLAST (call rates 99.9% vs 95.9%, respectively; p < 1×10^-15^), with significantly greater accuracy (false positive rates 0.46% vs 0.90%; p = 0.05). The heuristic information provided by SRST2 (that is, confident mismatches, insertions, deletions or truncations reported from read mapping) was a strong indicator of accuracy in the result: where an exact match was reported (98% of calls with depth ≥15x), the false positive rate was 0.2%; where an inexact match was reported, the false positive rate was 11.7%. For assembly, false positive rates were 0.2% for exact matches (95% of calls) and 76% for inexact matches. Hence, the key difference between the two methods was the ability of SRST2 to make correct calls where assembly+BLAST could not: for read depths ≥15x, SRST2 made a call with the correct allele 99.4% of the time, compared to only 95% for assembly analysis (p<1×10^-15^ for difference in rates). At sequence type (ST) level, the difference was even greater: SRST2 achieved accurate ST assignment for 96% of isolates with average depth ≥15x, whereas assembly+BLAST correctly identified only 76%.

**Figure 3.**
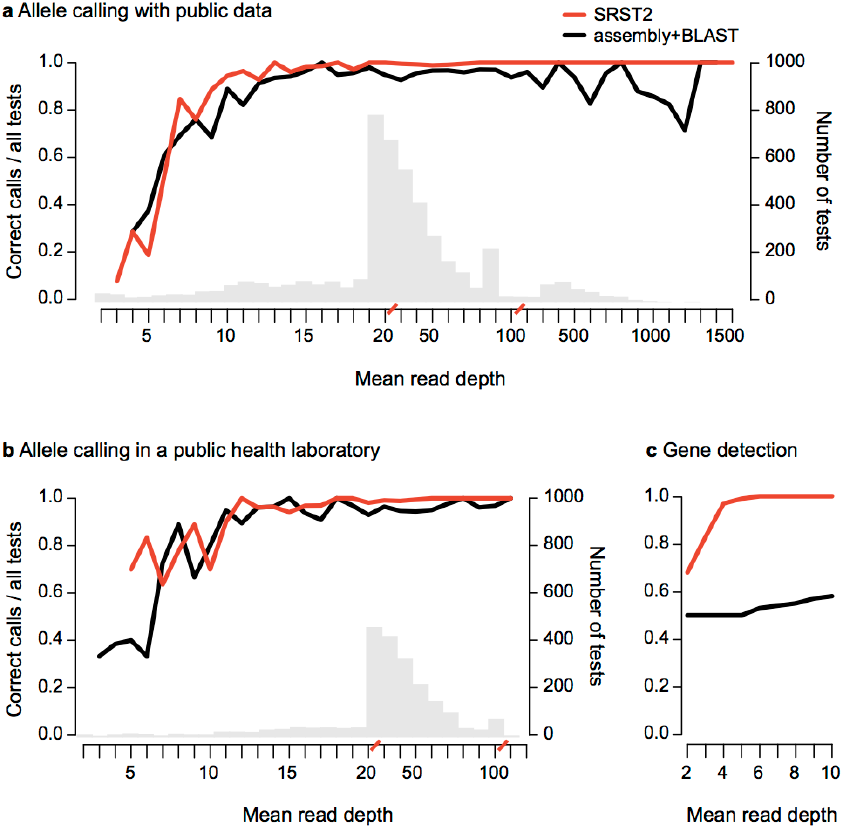
Overall accuracy of SRST2 allele calling and gene detection. (**a**) MLST analysis of public data from 5 species (N=543 genomes, 3801 loci, details Supplementary Table 1). Tests were grouped by read depth and accuracy rates (left y-axis, correct allele calls as a proportion of tests), calculated at each depth (x-axis, red slashes indicate scale change). Grey bars, number of tests at each depth (right y-axis); Lines, accuracy of allele calling. (**b**) MLST analysis of *Listeria monocytogenes* data (N=231 genomes, 1671 loci) conducted in a public health laboratory; colours and axes as in **a**. (**c**) Accuracy of *vanB* resistance gene detection for *E. faecium* read sets subsampled to low depth; y-axis shows proportion of correct (presence vs. absence) calls as a proportion of 100 tests at each depth; colours and axes as in **a**. A call of “present” implies detection of ≥90% of the length of the gene at ≥90% nucleotide identity.

To assess performance at low read depths (≤15x), ten *S. aureus* read sets were subsampled to low depths (**Methods**). This confirmed that an average depth of only 10x was required for SRST2 to achieve >90% call rate and <0.5% false positives (Figure 3a, Figure 4). MLST databases can be expected to grow indefinitely due to increasing diversity and broader sampling. However simulations (**Methods**) indicated that doubling the size of the *S. aureus* MLST database had no impact on SRST2 accuracy (Figure 3a, Figure 4).

**Figure 4.**
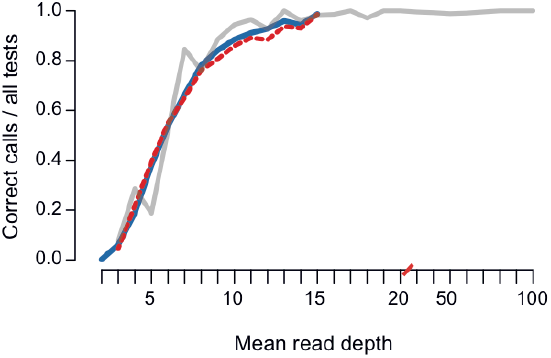
Accuracy of SRST2 allele calling at low read depths and with expanded MLST database size. MLST analysis of public *S. aureus* data. (N=10 read sets; each sampled 100 times to different depths; details in Methods). Tests were grouped by read depth and accuracy rates (y-axis, correct allele calls as a proportion of all tests), calculated at each depth (x-axis, red slashes indicate scale change from 1x to 10x). Red, real *S. aureus* MLST database; blue, expanded *S. aureus* MLST database (see Methods); grey, unsampled data from 5 species mapped to real databases (as shown in Figure 1, 3).

### Validation of gene detection using the vanA-B resistance gene

In addition to reliably distinguishing alleles of a given gene, SRST2 can also accurately determine the presence or absence of genes of interest, such as those encoding antimicrobial resistance or virulence. To evaluate this, we used 43 *E. faecium* genomes (Table 1), previously screened for vancomycin susceptibility and presence of the VanB vancomycin resistance operon *vanABHSXY*[35, 36]. Seventeen isolates were vancomycin resistant (VRE), and all were PCR positive for the *vanA-B* gene. These genomes were sequenced to ∼1,000x depth and SRST2 correctly detected *vanA-B* in 17/17 VRE. In five vancomycin sensitive (VSE) isolates PCR negative for *vanA-B*, SRST2 detected *VanA-B* sequences at very low depths (<0.2% of average depth), probably caused by minor but easily identifiable contamination during VRE-VSE multiplexed sequencing. SRST2 also confirmed the presence of the entire VanB operon, which is strongly predictive of the VRE phenotype. For comparison, assembly+BLAST identified full-length *vanA-B* sequences in just 7/17 VRE genomes, with multiple smaller hits spanning the full-length gene in five VRE and <50% coverage of the gene identified in the remaining five VRE. To investigate the effect of sequencing depth on gene detection, we randomly selected five VRE and five VSE read sets for subsampling at <10x average read depth. *VanA-B* was only ever detected in confirmed VRE genomes, and sensitivity of detection with SRST2 reached 100% for read sets with ≥5x average read depth (Fig. 1c).

To further explore the relative sensitivity of gene detection with SRST2, we screened all the read sets used for MLST validation (Table 1) for antimicrobial resistance genes in the ARG-Annot database of acquired resistance genes[18] (**Methods**). SRST2’s detection of whole genes was more sensitive than detection of whole or partial gene sequences by assembly+BLAST (Figure 5): 6.8% of genes detected at ≥90% coverage by SRST2 at depths ≥15x were not found at ≥90% coverage in assemblies. For most of these genes, smaller fragments were detected by BLAST (Figure 5); however, SRST2 has the advantage of sensitive detection and confident allele-calling across the full length of genes, even at low depths (Figure 3c, Figure 5).

**Figure 5.**
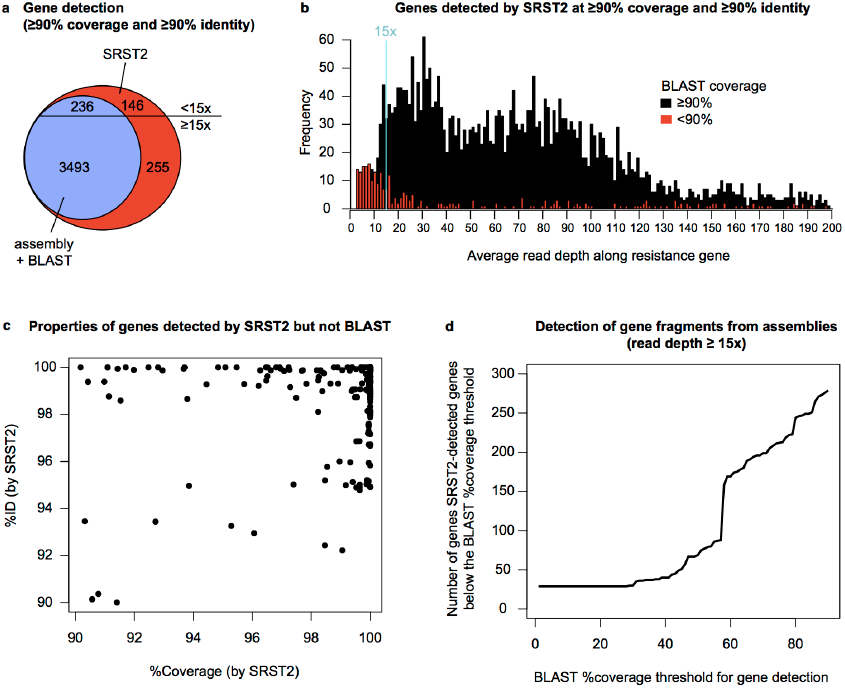
Resistance gene detection. (**a**) Venn diagram of antimicrobial resistance genes detected by SRST2 and assembly+BLAST, where the threshold for ‘detection’ of a gene is ≥90% coverage and ≥90% identity with a reference allele. No genes were detected by assembly+BLAST but not SRST2. (**b**) Distribution of average read depths per gene, calculated by SRST2 from mapped reads, for all genes detected by SRST2. (**c**) Coverage and nucleotide identity (%ID), as calculated by SRST2, for all genes detected by SRST2 but not by assembly+BLAST. (**d**) Impact of lowering the coverage threshold for detection of genes by BLAST (for those genes with ≥15x read depth).

### Validation of SRST2 in a public health laboratory

To validate SRST2 in a public health laboratory setting, we analysed 231 clinical isolates of *Listeria monocytogenes* and compared MLST data obtained from gold-standard PCR and amplicon sequencing with those obtained from SRST2 or assembly+BLAST analysis of Illumina MiSeq data (Figure 3b). Sequencing and analysis was performed by the Microbiological Diagnostic Unit Public Health Laboratory in Melbourne, Australia, the national reference laboratory for *L. monocytogenes*. For average read depths ≥15x, SRST2 had a substantially higher call rate than assembly-based analysis (99.6% vs. 95.7%; p < 1×10^-12^), with similar low false positive rates (0.7% vs. 0.6%; p=0.9). Hence, for samples with ≥15x data, a total of 99% of all alleles were called correctly by SRST2, a significantly higher proportion than the 95% achieved by assembly+BLAST (p<1×10^-12^). At <15x read depths, SRST2 also performed better than assembly-based analysis (87% vs 72% of alleles correctly called, respectively, p<1×10^-3^; Figure 3b).

Further, SRST2 is already being assessed for routine MLST analysis of *Streptococcus pneumoniae* at Public Health England (Anthony Underwood, personal communication), and the open-source SRST2 code has been adapted by Public Health Ontario, Canada to perform specialist *emm* typing of Group A *Streptococcus* [37].

## Demonstration of utility in hospital infection control

### Identification of antimicrobial resistant clones

In a hospital setting, the combination of MLST and gene detection can provide rapid and powerful insights for infection control without specialist bioinformatics knowledge. SRST2 analysis of 69 *K. pneumoniae* and 74 *E. coli* genomes from a UK hospital [38] revealed that each was dominated by a single ST with a high rate of the extended-spectrum beta-lactamase (ESBL) gene CTX-M-15 (*K. pneumoniae* ST490 comprising 25% of total, 71% of ESBL; *E. coli* ST131 comprising 40% of total, 77% of ESBL; Figure 6). Routine SRST2 surveillance of ESBL infections could be indicative of hospital outbreaks and used to identify which isolates should be investigated via transmission analysis.

**Figure 6.**
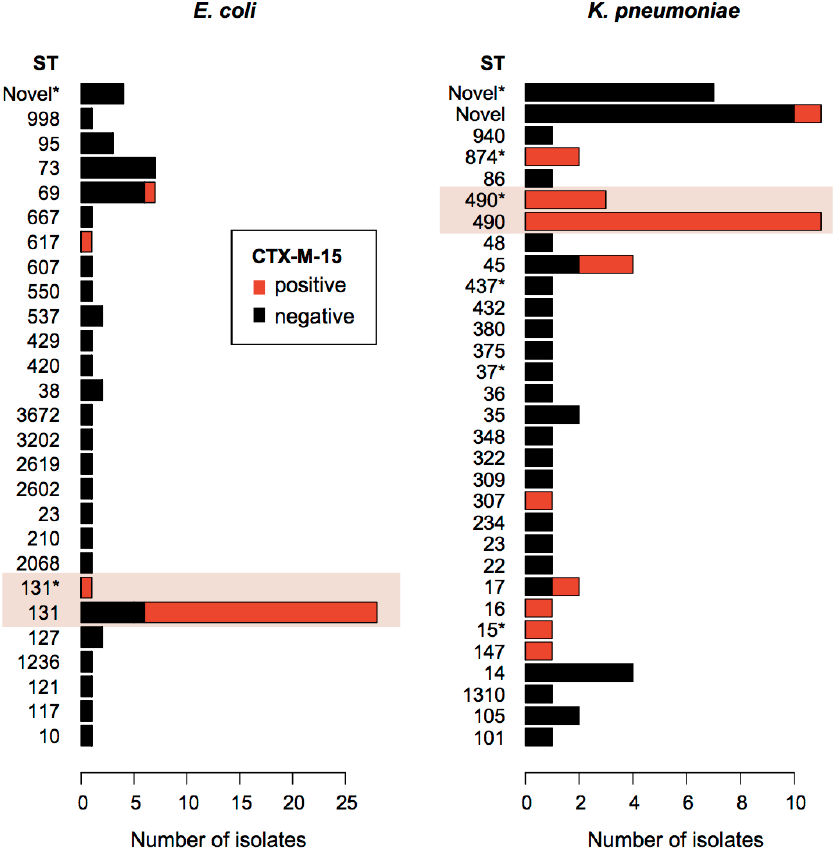
SRST2 analysis of sequence types and beta-lactamase CTX-M-15 genes amongst hospital isolates. Rates of isolation of different sequence types (STs), coloured by CTX-M-15 status, as determined by SRST2 run with default parameters on a public data set of strains from a single hospital. In each species, a single known ST dominates the population (highlighted) and is also the dominant source CTX-M-15 genes. ‘*’ next to an ST indicates a match to the closest defined ST; i.e., that for all 7 loci the closest known allele is the one belonging to that ST, however at ≥1 these loci there is an imprecise match (SNP or indel) compared to the known allele sequence. ‘Novel’ indicates a novel sequence type resulting from a combination of known alleles, with precise matches at all loci (‘NF’ in SRST2 output); ‘Novel*’ indicates a novel combination of alleles, with ≥1 of those alleles being novel itself (i.e. with no exact match in the MLST database) (‘NF*’ in SRST2 output).

Using the *E. faecium* genome data, collected as part of a 12-year hospital study of vancomycin resistance^25^, SRST2 took ∼30 minutes to generate the results and visualizations shown in Figure 7, indicating (i) increasing vancomycin resistance over time; (ii) a shift in dominant ST during the same period; and importantly (iii) that this was not attributable to the introduction nor transmission of a new resistant clone, as the resistance rates were steady (approximately 50%) across all dominant STs. Similar conclusions typically require many days of labour and specialised assays in the diagnostic laboratory [39] and have been confirmed by detailed WGS analysis showing frequent acquisition of VanB transposons by diverse circulating strains [35].

**Figure 7.**
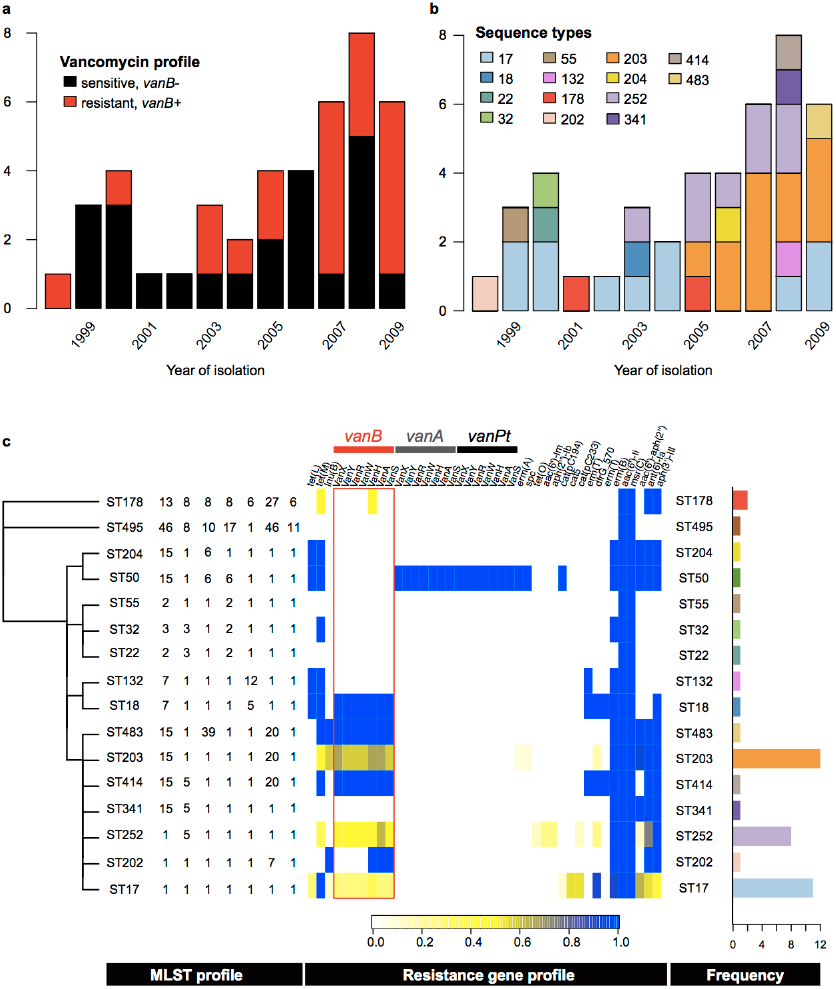
SRST2 analysis of *E. faecium* hospital data and hospital outbreak investigation. Temporal distribution of isolates is shown in (**a**) coloured by vancomycin resistance as inferred from *vanA-B* detection with SRST2, and in (**b**) by coloured by sequence type inferred by SRST2. (**c)** Summary of all SRST2 results by sequence type (ST), in order from left to right: single linkage clustering of STs by number of shared alleles; MLST allele profiles; heatmap indicating the proportion of isolates that carries each resistance gene (scale as indicated), frequency of the ST (axis as indicated, coloured as in **b**).

### Investigation of outbreaks and carbapenem resistance mechanisms

We next applied SRST2 to analyse data from real-world small-scale infection control investigations [15]. SRST2 took 5 minutes to generate results for suspected outbreaks of VRE and *E. cloaceae* (Figure 8), in which suspected outbreak isolates were readily distinguishable from epidemiologically unrelated isolates, consistent with WGS phylogenies and manual analysis of antimicrobial resistance markers [15]. SRST2 typing of 18 plasmid replicons [40] also indicated specific plasmid replicons (IncHI2, IncA/C) associated with two of the resistance profiles.

**Figure 8.**
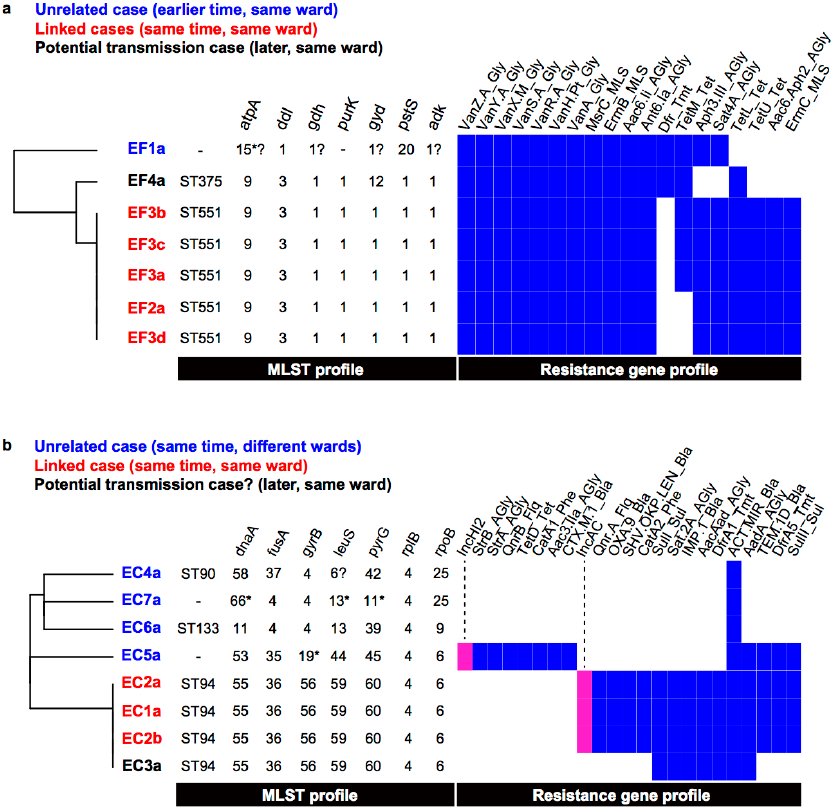
SRST2 analysis of hospital outbreak investigation. (**a**) Isolate genetic profiles obtained from SRST2 analysis, indicating that case EF4 was distinct in both sequence type and resistance gene profile from the outbreak cases EF2 and EF3. Full WGS analysis showed a similar result[15]. (**b**) Isolate genetic profiles obtained from SRST2 analysis, including plasmid replicons detected (pink). The profiles indicate that case EC3 shared the same sequence type as the linked cases EC1 and EC2 (ST94), but lacked the IncA/C plasmid and had a distinct resistance gene profile. Full WGS analysis showed that EC1 and EC2 isolates were much closer to each other (≤22 SNPs) than to EC3 (>150 SNPs)[15].

The authors also reported use of a complex hybrid of assembly, mapping and manual inspection to determine carbapenem resistance mechanisms in five Gram-negative bacteria isolated in close proximity [15]. SRST2 analysis of these five read sets identified the acquired beta-lactamases OXA-23 in AB223; IMP, SHV-12 and TEM-1 in EC1a; CTX-M-15 and TEM-1 in Eco216; CTX-M-15 and SHV-133 in KP652; and no acquired carbapenemase genes in EC302. These results are consistent with those reported from manual analysis [15].

## Methods

### Bacterial isolates and sequencing

A total of 231 *Listeria monocytogenes* isolates were analysed in this study, at the Microbiological Diagnostic Unit (MDU) Public Health Laboratory in Victoria, Australia. MDU is the national reference laboratory for *L. monocytogenes* and the isolates analysed include several from recent outbreaks as well as from the laboratory’s reference collection. Cultures of *L. monocytogenes* isolated from food, environmental or clinical specimens were purified by two successive single colony selections after streaking onto horse blood agar (HBA) incubated for 18-24 h at 37°C. Resultant bacterial growth on the surface of HBA medium was aseptically collected and resuspended in a cryotube (Nalgene) containing 1 mL of sterile glycerol storage broth (1.6% w/v Tryptone, Oxoid Pty Ltd, LP0042 containing 20% v/v glycerol) prior to storage at -70°C. Cultures were retrieved from storage as required and freshly grown (HBA, 18-24 h at 37°C) in preparation for DNA extraction. DNA was extracted from each isolate using QIAmp DNA Mini Kit (Qiagen) and eluted in EB buffer (Qiagen) (Tris buffer, no EDTA).

DNA samples were subjected to traditional *L. monocytogenes* MLST analysis [41, 42], with a minor modification to the annealing temperature for the *bglA* PCR (52°C not 45°C). The PCR products were purified with FastAP Thermosensitive Alkaline Phosphatase (Thermo Scientific) and Exonuclease I (Thermo Scientific). The purified PCR products were sequenced using BigDye Terminator v3 chemistry followed by capillary sequencing using a 3130xL Genetic Analyzer (Applied Biosystems). Trace analysis was conducted using BioNumerics version 6.6 with MLST Online plugin version 2.13 and Batch Sequence Assembly plugin version 1.34.

DNA was subjected to multiplex library preparation using Nextera XT followed by sequencing using an Illumina MiSeq. DNA was quantified by Qubit dsDNA HS Assay Kit (Invitrogen) and normalized to 0.2ng/µl. Total 1 ng of DNA was used for Nextera XT DNA Sample Preparation Kit (Illumina). Tagmentation of genomic DNA, PCR amplification with dual index primers, PCR clean-up using Agencourt AMPure XP (Beckman Coulter), DNA libraries normalization, library pooling and MiSeq sample loading were performed according to the manufacturer’s instruction with minor modifications. For longer than 2×250 bp runs on the MiSeq, 25 µl of AMPure XP beads was added to each PCR-amplified product during the PCR purification step otherwise 30 µl of AMPure XP beads was added. For some samples, after PCR purification, DNA fragment size and library concentration was analysed by 2100 Bioanalyzer (Agilent Technologies) and Qubit dsDNA HS Assay Kit (Invitrogen). DNA libraries were normalized manually to 4 nM and libraries with unique indexes were pooled in equal volumes. Each resulting pooled library was denatured and diluted with 0.2N NaOH and pre-chilled HT1 (Illumina) to produce a 20 pM denatured library in 1 mM NaOH. Prior to the MiSeq run, the denatured library was further diluted with pre-chilled HT1 to approximately 12-13.5 pM. 600µl of library including 2% (v/v) 20 pM denatured PhiX library (Illumina) was loaded together with MiSeq reagent kit v3 (Illumina) according to the manufacturer’s instructions.

### Publicly available short read data used in this study

Details of Illumina read sets used in this study are provided in Tables 1 and 2. Data tables specifying the expected STs of each read set, summarised from published papers, are available on the SRST2 website [31].

**Table 2.** Data sets used to demonstrate utility of SRST2 in the hospital setting

| Species | Citation | N | Average read depth | Read length |
| --- | --- | --- | --- | --- |
| *Enterococcus faecium* (Fig 2a-c) | [35] | 43 | 658 | 101 |
| Hospital outbreak investigations (Fig 2d-e) | [15] | 20 | 36x | 151 |
| *K. pneumoniae, E. coli* | [38] | 69, 74 | 34x | 101 |

### Subsampling of read sets

To explore accuracy at low read depths, ten genomes each of *S. aureus* and *E. faecium* were selected for random subsampling of reads to simulate genomes sequenced to low read depth. To do this, we used the mean read depth across MLST loci to calculate the sampling fraction required to achieve approximately 1x, 2x, … 10x mean read depth. We randomly sampled reads from the forward reads file at the required sampling fraction, and extracted the corresponding reverse reads, using Perl scripts. Ten random samples were generated from each read set at each depth level, generating a total of 1,000 read sets for each species.

### Sequence databases used in this study

MLST databases for *Staphylococcus aureus*, *Streptococcus pneumoniae*, *Salmonella enterica*, *Escherichia coli*, *Enterococcus faecium*, *Listeria monocytogenes* and *Enterobacter cloaceae* were downloaded from pubmlst.org using the *getmlst.py* script included with SRST2 (June 2014).

Antimicrobial resistance gene detection was performed using the ARG-Annot database of acquired resistance genes [18]. Allele sequences (DNA) were downloaded in fasta format[43] (May, 2014). Sequences were clustered into gene groups with ≥80% identity using CD-hit[44] and the headers formatted for use with SRST2 using the scripts provided (*cdhit_to_csv.py*, *csv_to_gene_db.py*). A copy of the formatted sequence database used in this study is included in the SRST2 github repository [31].

Representative sequences for 18 plasmid replicons were extracted from GenBank using the accessions and primer sequences specified by Carattoli *et al* [40]. A copy of the formatted sequence database used in this study is included in the SRST2 github repository [31].

### Simulation of expanded S. aureus MLST database

As more genomes are sequenced and as bacteria continue to evolve, novel alleles will continue to be discovered and thus the size of allele databases will increase. To explore the impact of database size on accuracy of allele detection with SRST2, we simulated expansion of the current *S. aureus* MLST database from 2,161 alleles (mean 309 per locus) to 5,578 alleles (mean 797 per locus). The additional ∼500 alleles per locus were generated using *netrecodon* v6.0.0 [45]. Sequences derived from the true MLST database were used to seed the simulation at each locus as follows. Existing alleles were translation-aligned between start (alignment start) and stop (alignment end) codons, those containing a frameshift or stop codon were removed, and the modal consensus sequence was exported. The best-fit DNA substitution model of each true alignment was determined using the AIC in *MrModeltest* v2.3, as implemented in *PAUP\** v4.0b. In *netrecodon*, the modal sequences were forward evolved under the coalescent, using the parameters of the best-fitting model for each locus, mutation rate 1E-7 and recombination rate 1E-7/15 (based on reported r/m of 1/15 [46]). A total of 100 independent replicates of forward evolution were performed per locus, retaining 2,000 sequences per replicate (N = 200,000 simulated sequences per locus). The first 500 unique simulated sequences at each locus were added to the MLST database, and duplicate sequences were removed.

### Analysis runs and time calculations

All SRST2, assembly and BLAST analysis was run on a Linux cluster (iDataplex x86 system, “Barcoo” cluster at VLSCI [47]). SRST2 was run with default parameters. Details of *Velvet* assembly and BLAST analysis are given below. Run times were calculated from time stamps extracted from log files for SRST2 and *Velvet Optimiser* assembly runs.

### Assembly-based analysis

Assemblies were generated using the de novo assembler *Velvet* v1.2.10 [24], with optimal kmer choice for each readset refined through iterative calls to *VelvetOptimiser* v2.2.5 [34]. Briefly, each read set was assembled using a call to *VelvetOptimiser* with kmers from 29 up to 89, in steps of 12. The optimal kmer, k_1_, was extracted and a second call to *VelvetOptimiser* was made using kmers from k_1_-12 up to k_1_+12, in steps of 4. A final call to *VelvetOptimiser* was run using kimers from k_2_-4 up to k_2_+4, in steps of 2. The final assembly was that output from the third and final call to *VelvetOptimiser*.

For MLST analysis from assemblies, a nucleotide BLAST+ (v2.2.25) search was performed for each locus and each contig set. In this BLAST search, the contig set was used to query the database containing all known allele sequences for a given locus, and the top BLAST hit was reported. If this hit had ≥90% nucleotide identity across ≥90% of the length of the reference allele sequence, an allele call was recorded. If the hit was an exact match to a known allele (i.e. 100% nucleotide identity across 100% of the length of the allele sequence), this was considered a precise allele call. The Python code used is available within the SRST2 distribution. For gene detection analysis from assemblies, a nucleotide BLAST search was performed in which the set of reference sequences (sequence database, i.e. antimicrobial resistance gene database) was used to query the database of all contigs for that assembly.

### Statistical analysis

All statistical analysis and data plotting was performed in *R*. Allele calling performance of SRST2 and assembly+BLAST was assessed via three metrics: (i) call rate = total number of allele calls made, for SRST2 this was a call with ≥90% coverage and no uncertainty recorded (i.e. with ≥2x read depth at both ends and also neighbouring any truncations or deleted bases), for BLAST this was a call with ≥90% coverage and ≥90% nucleotide identity; (ii) false positive rate = total number of correct allele calls as a proportion of all calls; (iii) proportion of all tests resulting in a call with a correct allele, equal to (call rate) * [1 – (false positive rate)]. As these metrics are proportions, the significance of differences in performance metrics was calculated using a two-sided test for equality of proportions (*prop.test* function in *R*). Resistance gene detection was assessed using a cut-off of ≥90% coverage and ≥90% identity to define the presence of a gene.

## Acknowledgments

This work was supported by the NHMRC of Australia (grant #1043830; fellowships #1061409 (KEH) and #1061435 (MI, co-funded with the Australian Heart Foundation)) and the Victorian Life Sciences Computation Initiative (VLSCI) (grant #VR0082).

## Author Contributions

Wrote code: MI, HD, BJP, KEH. Designed the study and algorithm: MI, BJP, JZ, KEH. Performed DNA extraction and sequencing: TT. Analyzed data: MI, HD, LR, MBS, KEH. All authors read and approved the final manuscript.

## Additional data files

The following additional data are available with the online version of this paper. Additional data file 1 shows separate plots for call rates and true positive rates for the six public data sets used for MLST allele typing validation (these two measures were combined to give the overall accuracy plot in Figure 3). Additional data file 2 is a CSV table listing the 543 read file accessions from these data sets together with the corresponding expected sequence types (STs), which were extracted from published results of PCR and capillary sequencing and used to assess accuracy of SRST2 allele calling (shown in Figure 1a).

## References

1. Sabat AJ, Budimir A, Nashev D, Sa-Leao R, van Dijl J, Laurent F, Grundmann H, Friedrich AW, Markers ESGoE: Overview of molecular typing methods for outbreak detection and epidemiological surveillance. Euro Surveill 2013, 18:20380.

2. Bertelli C, Greub G: Rapid bacterial genome sequencing: methods and applications in clinical microbiology. Clin Microbiol Infect 2013, 19:803–813.

3. Maiden MC: Multilocus sequence typing of bacteria. Annu Rev Microbiol 2006, 60:561–588.

4. Gilmour MW, Graham M, Reimer A, Van Domselaar G: Public health genomics and the new molecular epidemiology of bacterial pathogens. Public Health Genomics 2013, 16:25–30.

5. Pallen MJ, Loman NJ, Penn CW: High-throughput sequencing and clinical microbiology: progress, opportunities and challenges. Curr Opin Microbiol 2010, 13:625–631.

6. Joseph SJ, Read TD: Bacterial population genomics and infectious disease diagnostics. Trends Biotechnol 2010, 28:611–618.

7. Didelot X, Bowden R, Wilson DJ, Peto TE, Crook DW: Transforming clinical microbiology with bacterial genome sequencing. Nat Rev Genet 2012, 13:601–612.

8. Didelot X, Eyre DW, Cule M, Ip CL, Ansari MA, Griffiths D, Vaughan A, O’Connor L, Golubchik T, Batty EM, et al: Microevolutionary analysis of Clostridium difficile genomes to investigate transmission. Genome Biol 2012, 13:R118.

9. Price JR, Didelot X, Crook DW, Llewelyn MJ, Paul J: Whole genome sequencing in the prevention and control of Staphylococcus aureus infection. J Hosp Infect 2013, 83:14–21.

10. Koser CU, Ellington MJ, Cartwright EJ, Gillespie SH, Brown NM, Farrington M, Holden MT, Dougan G, Bentley SD, Parkhill J, Peacock SJ: Routine use of microbial whole genome sequencing in diagnostic and public health microbiology. PLoS Pathog 2012, 8:e1002824.

11. Rohde H, Qin J, Cui Y, Li D, Loman NJ, Hentschke M, Chen W, Pu F, Peng Y, Li J, et al: Open-source genomic analysis of Shiga-toxin-producing E. coli O104:H4. New Engl J Med 2011, 365:718–724.

12. Eyre DW, Golubchik T, Gordon NC, Bowden R, Piazza P, Batty EM, Ip CL, Wilson DJ, Didelot X, O’Connor L, et al: A pilot study of rapid benchtop sequencing of Staphylococcus aureus and Clostridium difficile for outbreak detection and surveillance. BMJ Open 2012, 2.

13. Torok ME, Peacock SJ: Rapid whole-genome sequencing of bacterial pathogens in the clinical microbiology laboratory--pipe dream or reality? J Antimicrob Chemother 2012, 67:2307–2308.

14. Harris SR, Cartwright EJ, Torok ME, Holden MT, Brown NM, Ogilvy-Stuart AL, Ellington MJ, Quail MA, Bentley SD, Parkhill J, Peacock SJ: Whole-genome sequencing for analysis of an outbreak of meticillin-resistant Staphylococcus aureus: a descriptive study. The Lancet infectious diseases 2012.

15. Reuter S, Ellington MJ, Cartwright EJ, Koser CU, Torok ME, Gouliouris T, Harris SR, Brown NM, Holden MT, Quail M, et al: Rapid bacterial whole-genome sequencing to enhance diagnostic and public health microbiology. JAMA Intern Med 2013, 173:1397–1404.

16. Sherry NL, Porter JL, Seemann T, Watkins A, Stinear TP, Howden BP: Outbreak investigation using high-throughput genome sequencing within a diagnostic microbiology laboratory. J Clin Microbiol 2013, 51:1396–1401.

17. D’Auria G, Schneider MV, Moya A: Live Genomics for Pathogen Monitoring in Public Health. Pathogens 2014, 3:93–108.

18. Gupta SK, Padmanabhan BR, Diene SM, Lopez-Rojas R, Kempf M, Landraud L, Rolain JM: ARG-ANNOT, a new bioinformatic tool to discover antibiotic resistance genes in bacterial genomes. Antimicrob Agents Chemother 2014, 58:212–220.

19. Zankari E, Hasman H, Cosentino S, Vestergaard M, Rasmussen S, Lund O, Aarestrup FM, Larsen MV: Identification of acquired antimicrobial resistance genes. J Antimicrob Chemother 2012, 67:2640–2644.

20. Carattoli A, Zankari E, Garcia-Fernandez A, Voldby Larsen M, Lund O, Villa L, Moller Aarestrup F, Hasman H: In Silico Detection and Typing of Plasmids using PlasmidFinder and Plasmid Multilocus Sequence Typing. Antimicrob Agents Chemother 2014, 58:3895–3903.

21. Larsen MV, Cosentino S, Rasmussen S, Friis C, Hasman H, Marvig RL, Jelsbak L, Ponten TS, Ussery DW, Aarestrup FM, Lund O: Multilocus Sequence Typing of Total Genome Sequenced Bacteria. J Clin Microbiol 2012.

22. Jolley KA, Maiden MC: Automated extraction of typing information for bacterial pathogens from whole genome sequence data: Neisseria meningitidis as an exemplar. Euro Surveill 2013, 18:20379.

23. Cody AJ, McCarthy ND, Jansen van Rensburg M, Isinkaye T, Bentley SD, Parkhill J, Dingle KE, Bowler IC, Jolley KA, Maiden MC: Real-time genomic epidemiological evaluation of human Campylobacter isolates by use of whole-genome multilocus sequence typing. J Clin Microbiol 2013, 51:2526–2534.

24. Zerbino DR, Birney E: Velvet: algorithms for de novo short read assembly using de Bruijn graphs. Genome Res 2008, 18:821–829.

25. Bankevich A, Nurk S, Antipov D, Gurevich AA, Dvorkin M, Kulikov AS, Lesin VM, Nikolenko SI, Pham S, Prjibelski AD, et al: SPAdes: a new genome assembly algorithm and its applications to single-cell sequencing. J Comput Biol 2012, 19:455–477.

26. Koren S, Treangen TJ, Hill CM, Pop M, Phillippy AM: Automated ensemble assembly and validation of microbial genomes. BMC Bioinformatics 2014, 15:126.

27. Salzberg SL, Phillippy AM, Zimin A, Puiu D, Magoc T, Koren S, Treangen TJ, Schatz MC, Delcher AL, Roberts M, et al: GAGE: A critical evaluation of genome assemblies and assembly algorithms. Genome research 2012, 22:557–567.

28. Inouye M, Conway TC, Zobel J, Holt KE: Short read sequence typing (SRST): multi-locus sequence types from short reads. BMC Genomics 2012, 13:338.

29. Loman NJ, Constantinidou C, Chan JZ, Halachev M, Sergeant M, Penn CW, Robinson ER, Pallen MJ: High-throughput bacterial genome sequencing: an embarrassment of choice, a world of opportunity. Nat Rev Microbiol 2012, 10:599–606.

30. Meacham F, Boffelli D, Dhahbi J, Martin DI, Singer M, Pachter L: Identification and correction of systematic error in high-throughput sequence data. BMC Bioinformatics 2011, 12:451.

31. SRST2 - Short Read Sequence Typing for Bacterial Pathogens [http://katholt.github.io/srst2/]

32. Langmead B, Salzberg SL: Fast gapped-read alignment with Bowtie 2. Nat Methods 2012, 9:357–359.

33. Li H, Handsaker B, Wysoker A, Fennell T, Ruan J, Homer N, Marth G, Abecasis G, Durbin R: The Sequence Alignment/Map format and SAMtools. Bioinformatics 2009, 25:2078–2079.

34. VelvetOptimiser [http://bioinformatics.net.au/software.velvetoptimiser.shtml]

35. Howden BP, Holt KE, Lam MM, Seemann T, Ballard S, Coombs GW, Tong SY, Grayson ML, Johnson PD, Stinear TP: Genomic insights to control the emergence of vancomycin-resistant enterococci. MBio 2013, 4.

36. Stinear TP, Olden DC, Johnson PD, Davies JK, Grayson ML: Enterococcal vanB resistance locus in anaerobic bacteria in human faeces. Lancet 2001, 357:855–856.

37. Athey TB, Teatero S, Li A, Marchand-Austin A, Beall BW, Fittipaldi N: Deriving Group A Streptococcus Typing Information from Short-Read Whole Genome Sequencing Data. J Clin Microbiol 2014.

38. Stoesser N, Batty EM, Eyre DW, Morgan M, Wyllie DH, Del Ojo Elias C, Johnson JR, Walker AS, Peto TE, Crook DW: Predicting antimicrobial susceptibilities for Escherichia coli and Klebsiella pneumoniae isolates using whole genomic sequence data. J Antimicrob Chemother 2013, 68:2234–2244.

39. Johnson PD, Ballard SA, Grabsch EA, Stinear TP, Seemann T, Young HL, Grayson ML, Howden BP: A sustained hospital outbreak of vancomycin-resistant Enterococcus faecium bacteremia due to emergence of vanB E. faecium sequence type 203. J Infect Dis 2010, 202:1278–1286.

40. Carattoli A, Bertini A, Villa L, Falbo V, Hopkins KL, Threlfall EJ: Identification of plasmids by PCR-based replicon typing. J Microbiol Methods 2005, 63:219–228.

41. Ragon M, Wirth T, Hollandt F, Lavenir R, Lecuit M, Le Monnier A, Brisse S: A new perspective on Listeria monocytogenes evolution. PLoS Pathog 2008, 4:e1000146.

42. Listeria monocytogenes MLST Database [http://www.pasteur.fr/recherche/genopole/PF8/mlst/Lmono.html]

43. ARG-ANNOT - Antibiotic Resistance Gene-ANNOTation [http://www.mediterranee-infection.com/article.php?laref=282&titer=arg-annot]

44. Li W, Godzik A: Cd-hit: a fast program for clustering and comparing large sets of protein or nucleotide sequences. Bioinformatics 2006, 22:1658–1659.

45. Arenas M, Posada D: Coalescent simulation of intracodon recombination. Genetics 2010, 184:429–437.

46. Feil EJ, Cooper JE, Grundmann H, Robinson DA, Enright MC, Berendt T, Peacock SJ, Smith JM, Murphy M, Spratt BG, et al: How clonal is Staphylococcus aureus? J Bacteriol 2003, 185:3307–3316.

47. VIctorian Life Sciences Computation Initiative [http://www.vlsci.org.au/]

48. Holden MT, Hsu LY, Kurt K, Weinert LA, Mather AE, Harris SR, Strommenger B, Layer F, Witte W, de Lencastre H, et al: A genomic portrait of the emergence, evolution, and global spread of a methicillin-resistant Staphylococcus aureus pandemic. Genome Res 2013, 23:653–664.

49. Castillo-Ramirez S, Corander J, Marttinen P, Aldeljawi M, Hanage WP, Westh H, Boye K, Gulay Z, Bentley SD, Parkhill J, et al: Phylogeographic variation in recombination rates within a global clone of methicillin-resistant Staphylococcus aureus. Genome Biol 2012, 13:R126.

50. Croucher NJ, Harris SR, Fraser C, Quail MA, Burton J, van der Linden M, McGee L, von Gottberg A, Song JH, Ko KS, et al: Rapid pneumococcal evolution in response to clinical interventions. Science 2011, 331:430–434.

51. Okoro CK, Kingsley RA, Connor TR, Harris SR, Parry CM, Al-Mashhadani MN, Kariuki S, Msefula CL, Gordon MA, de Pinna E, et al: Intracontinental spread of human invasive Salmonella Typhimurium pathovariants in sub-Saharan Africa. Nat Genet 2012, 44:1215–1221.

52. Holt K, Baker S, Weill F, Holmes E, Kitchen A, Yu J, Sangal V, Brown D, Coia J, Kim D, et al: Shigella sonnei genome sequencing and phylogenetic analysis indicate recent global dissemination from Europe. Nat Genet 2012, 44:1056–1059.

